# Ribosome profiling analysis of eEF3-depleted *Saccharomyces cerevisiae*

**DOI:** 10.1101/413252

**Authors:** Villu Kasari, Tõnu Margus, Gemma C. Atkinson, Marcus J.O. Johansson, Vasili Hauryliuk

## Abstract

In addition to the standard set of translation factors common in eukaryotic organisms, protein synthesis in the yeast *Saccharomyces cerevisiae* requires an ABCF ATPase factor eEF3, eukaryotic Elongation Factor 3. eEF3 is an E-site binder that was originally identified as an essential factor involved in the elongation stage of protein synthesis. Recent biochemical experiments suggest an additional function of eEF3 in ribosome recycling. We have characterised the global effects of eEF3 depletion on translation using ribosome profiling. Depletion of eEF3 results in decreased ribosome density at the stop codon, indicating that ribosome recycling does not become rate limiting when eEF3 levels are low. Consistent with a defect in translation elongation, eEF3 depletion causes a moderate redistribution of ribosomes towards the 5’ part of the open reading frames. We observed no E-site codon-or amino acid-specific ribosome stalling upon eEF3 depletion, supporting its role as a general elongation factor. Surprisingly, depletion of eEF3 leads to a relative decrease in P-site proline stalling, which we hypothesise is a secondary effect of generally decreased translation and/or decreased competition for the E-site with eIF5A.

## Introduction

Protein synthesis – translation – is universally performed by the ribosome, which is assisted by specialised proteins referred to as translation factors. Some translation factors are universally conserved, e.g. the elongation factor eEF2/EF-G^1^– and some are lineage-specific, such as elongation factor 3, eEF3, a member of the ABCF ATPase family^2,3^. While initial analysis of eEF3 distribution suggested it a fungi-specific translational factor^4^, its distribution is broader, with eEF3-like homologues found in non-fungal species, such as oomycete *Phytophthora infestans*^5^, choanoflagellates, and various distantly related algae^3^. The protein is essential both for the viability of *Saccharomyces cerevisiae*^6^ and for peptide elongation in a reconstituted yeast translational system^7,8^. Despite decades of research it is not clear why eEF3 is essential for translation elongation in yeast, since it is not a part of the translational apparatus in the vast majority of eukaryotes, including animals and land plants3. In the test tube eEF3 stimulates aminoacyl-tRNA delivery by elongation factor eEF1A^8,9^, with the C-terminal region of eEF3 directly interacting with eEF1A^10,11^. In addition to elongation, biochemical experiments suggest a secondary function for eEF3 in ribosome recycling^12^, inviting an analogy with the multifunctional bacterial GTPase EF-G that participates both in elongation^13^and ribosome recycling^14^. In a reconstituted biochemical system, eEF3-mediated ribosome recycling does not lead to an accumulation of ribosome ‘halfmers’ – that is 40S subunits associated with mRNA after the 60S release^12,15^ Therefore, Kurata and colleagues proposed that eEF3 mediates a recycling pathway via ejection of P-site tRNA and mRNA from the 80S termination complex. Cryoelectron microscopy reconstruction of ribosome-associated eEF3 localises the factor in the vicinity of the ribosomal E-site^16^, providing a structural explanation for the biochemical observation that eEF3 competes with the E-site tRNA on the ribosome in the presence of ATP^17^. We have analysed the global effects of eEF3 depletion on translation in *S. cerevisiae* using ribosome profiling (Ribo-Seq), a functional genomics approach that provides a bird’s-eye view of mRNA translation in the cell by means of isolation and sequencing of mRNA fragments protected by translating ribosomes^18^-^21^. We took advantage of our high-coverage dataset to ask the following questions: which stage of the ribosomal functional cycle is more sensitive to eEF3 depletion – elongation or ribosome recycling? And is eEF3’s function in elongation codon-or amino acid-specific?

## Results

### Construction and characterisation of the *P*_MET25_-YEF3 strain for tunable eEF3 expression

To investigate the role of eEF3 in translation, we set out to develop a system that would allow quick, specific and efficient depletion of eEF3. As our first approach, we constructed a set of strains in which the synthesis of different destabilised forms of eEF3 is post-transcriptionally inhibited by addition of tetracycline to the medium^22^(see Supplementary information S1). However, even in the case of the most responsive strain, eEF3 depletion was inefficient and did not cause growth inhibition until after 7-8 hours. Therefore, rather than rely on rapid eEF3 depletion, we opted for controlling the steady-state level of the protein. We constructed a strain in which the sequence upstream of the endogenous *YEF3* ORF, encoding eEF3, was replaced with the sequence of the promoter and 5′-UTR of the methionine-repressible *MET25* gene^23^. The resulting P*_MET25_*-*YEF3* strain shows concentration-dependent growth inhibition upon addition of methionine to liquid (Fig. 1a) and solid (Fig. 1b) medium. In good agreement with the growth assays, western blotting revealed that the abundance of eEF3 decreases with increasing concentration of methionine (Fig. 1c and Supplementary Fig. S1). Moreover, the eEF3 levels in the P**_MET25_*-YEF3* strain grown in the absence of methionine are comparable to those in wild-type cells. Importantly, the levels of eukaryotic elongation factor 2 (eEF2), ribosomal proteins Rps8 and Rps10 as well as phosphoglycerate kinase 1 (Pgk1), are largely unaffected by the methionine concentration in the medium, demonstrating the specificity of eEF3 depletion.

**Figure 1.**
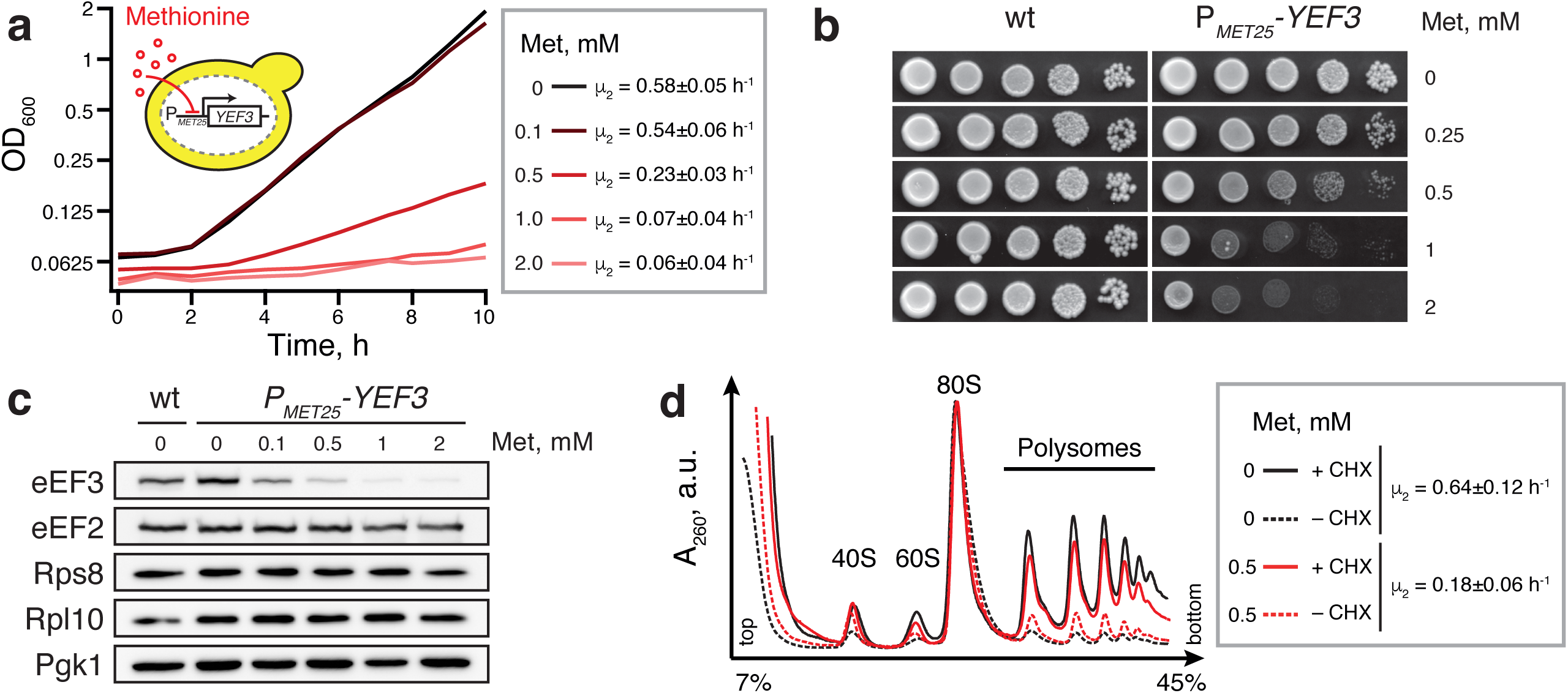
Tunable repression of eEF3 expression leads to a gradual decrease in growth rate. (**a**) The P*_MET25_*-*YEF3* (VKY8) strain was grown at 30°C in liquid synthetic complete medium lacking methionine and cysteine (SC-met-cys) supplemented with methionine at different concentrations (see insert). The growth rate (µ _2_) was calculated as the slope of the linear regression of log_2_-transformed OD_600_ measurements. (**b**) Wild type (VKY9) and P*_MET25_*-*YEF3* (VKY8) strains were grown overnight in SC-met-cys medium, 10-fold serially diluted, spotted on SC-met-cys plates supplemented with the indicated concentration of methionine, and incubated at 30°C for two days. (**c**) Western blot analysis of the P*_MET25_*-*YEF3* strain grown in SC-met-cys medium supplemented with indicated methionine concentrations. In addition to eEF3, the blot was probed for eEF2, Pgk1, Rps8 and Rpl10. Full-length western blots are presented in Supplementary Fig. S1. (**d**) Polysome profile analyses of the P*_MET25_*-*YEF3* strain grown in the presence and absence of 0.5 mM methionine. Before harvesting, the cells were either treated with 100 µg/ml cycloheximide for 10 minutes (+CHX) or left untreated (–CHX, ‘run-off conditions’). Whole cell extracts were resolved on sucrose gradients and the absorbance at 260 nm was measured during fractionation.

### Depletion of eEF3 decreases growth without dramatically perturbing translation

To assess the overall effect of eEF3 depletion on translation, we used velocity sedimentation of whole cell lysates in sucrose gradients to analyze the polysome profiles of the P_*MET25*_-*YEF3* strain grown in the absence or presence of 0.5 mM methionine. In these experiments, the presence of 0.5 mM methionine in the medium increased the generation time from 1.6 to 5.6 hours and decreased the eEF3 levels at least five times (Supplementary Fig. S2). Prior to the preparation of the lysates, the cultures were either pre-treated for 10 minutes with 100 µg/ml cycloheximide, CHX, in order to stabilise polysomes, or left untreated (‘run-off conditions’) (Fig. 1d). Depletion of eEF3 results in a slight decrease of the polysomal fraction in the presence of CHX (compare black and red solid lines). However, in run-off conditions the polysomal fraction is slightly larger in eEF3 depleted cells (black and red dashed lines). The effect of eEF3 depletion on polysome run-off is consistent with the previous finding that polysomes are stabilised in cells with a mutant form of eEF3^11^. While reduced polysome run-off is likely a consequence of a defect in translation elongation, it can, in principle, also reflect a ribosome recycling defect, i.e., the queueing of elongating ribosomes behind ribosomes stalled at stop codons.

### Generation and technical analysis of Ribo-Seq and RNA-Seq datasets

Ribo-Seq analysis allows sensitive identification of specific rate-limiting steps in translation manifested in an increased read density of ribosome protected fragments (RPFs). To uncover the specific effects of eEF3 depletion, we applied ribosome profiling to the P*_MET25_*-*YEF3* strain growing exponentially in the presence or absence of 0.5 mM methionine. The yeast cultures used for Ribo-Seq were not treated with CHX prior to harvesting since CHX treatment can cause a systematic bias in *S. cerevisiae* ribosome profiling experiments^24,25^. However, CHX was added in the lysis buffer to inhibit translation elongation and avoid ‘run-off’ during purification of the ribosome protected RNA fragments. Ribo-Seq and RNA-Seq libraries were prepared and sequenced from two biological replicates. Ribo-Seq (25-35 nt) and RNA-Seq (50 nt) reads were mapped to the genome allowing only unique alignments. The detailed description of the NGS data analysis pipeline is described in the Methods section as well as at the GitHub depository page (https://github.com/GCA-VH-lab/RiboSeqPy).

Prior to the analysis of the specific effects of eEF3 depletion on translation, we scrutinised the specificity and reproducibility of our Ribo-Seq and RNA-Seq datasets. In good agreement with the western blotting results, growth in medium containing 0.5 mM methionine dramatically reduced the number of ribosome footprints mapped to the *YEF3* ORF (Fig. 2a). Moreover, the ribosome occupancy in the PGK1 ORF is comparable between the two growth conditions (Fig. 2b). The two biological Ribo-Seq and RNA-Seq replicates demonstrate high reproducibility. The R^2^for the Ribo-Seq data, quantified in reads per million (RPM) for individual ORFs (Ribo-Seq^ORF^), is 0.977 in the presence, and 0.993 in the absence of methionine (Fig. 2c and Supplementary Fig. S3a). The RNA-Seq^ORF^ data also show excellent reproducibility between the two replicates (Fig. 2d and Supplementary Fig. S3b, R^2^=0.931 and R^2^=0.949). To uncover codon-specific effects in the ribosome profiling datasets, we used individual 5’ offsets (12-15 nucleotides) to assign the position of the ribosomal P-site in the 28-35 nucleotide (nt) reads (Supplementary Fig. S4 and S5). The reproducibility of the Ribo-Seq libraries quantified per individual codon (Ribo-Seq^codon^) is still good but lower than that quantified per ORF (Fig. 2e and Supplementary Fig. S3c, R^2^ =0.852 and R^2^ =0.803). To examine effects on translation efficiency, we also calculated the ribosomal load, defined as the ratio between the Ribo-Seq and RNA-Seq coverage. The ribosomal load was calculated either per individual ORF (ribosomal load_ORF_, used for differential gene expression analyses, Fig. 3c, see below) or per individual nucleotide position (ribosomal load_position_, used for metagene and polarity score plots, Fig. 4, see below). The ribosomal load is highly reproducible when calculated per ORF (R^2^_ribosomal load ORF 0 mM Met_ =0.932, Supplementary Fig. S3d and R^2^_ribosomal load ORF 0 mM Met_ 0.889, Supplementary Fig. S3e).

**Figure 2.**
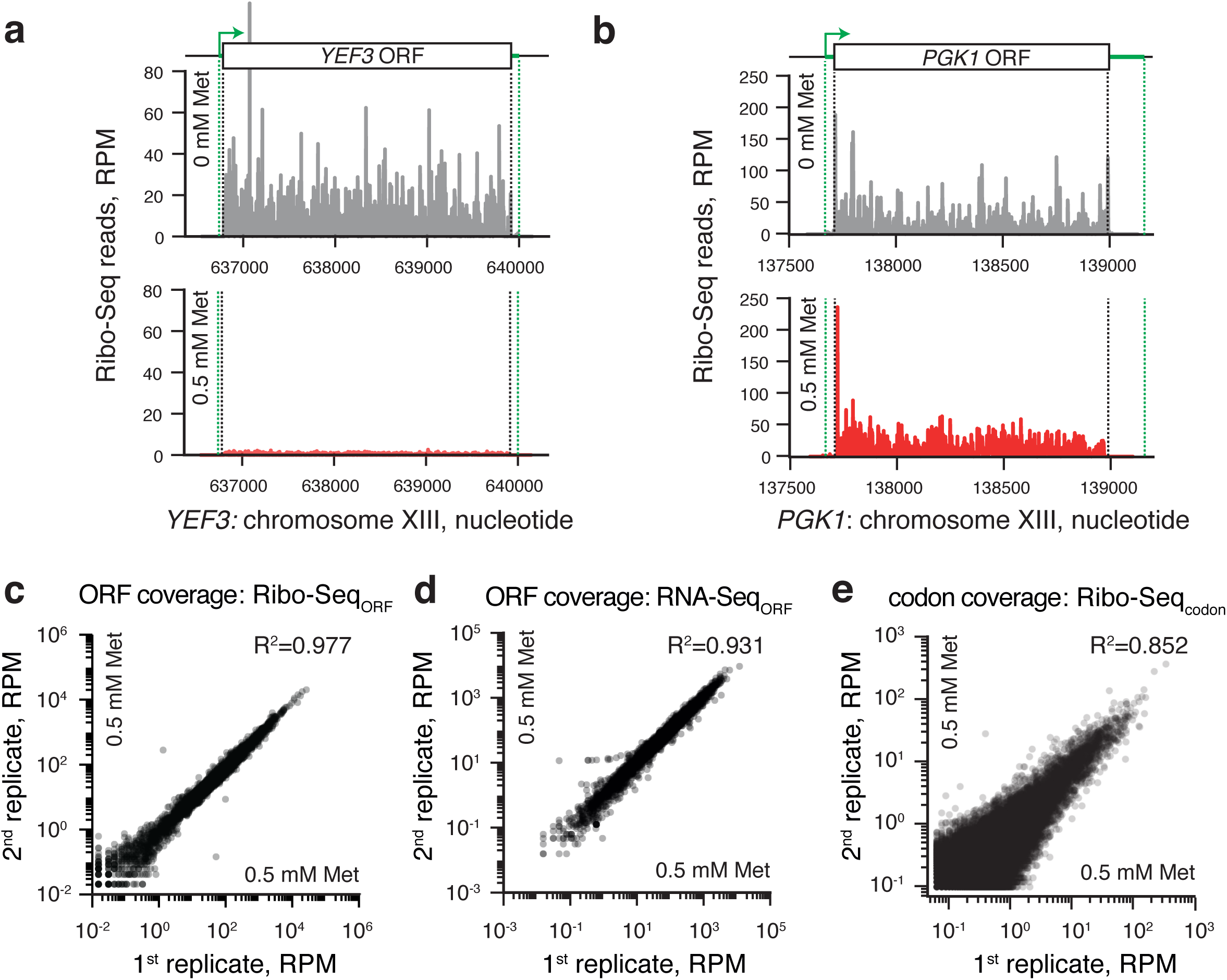
Reproducibility and specificity of Ribo-Seq and RNA-Seq datasets. Ribosome footprint density along the *YEF3* (**a**) and PGK1 (**b**) genes in the absence of methionine (upper panel, grey trace, no repression of eEF3 expression) and presence of 0.5 mM methionine (lower panel, red trace, repression of eEF3 expression). Green dotted lines indicate mRNA 3’ and 5’ ends. Reproducibility of ribosome footprint (**c** and **e**) and RNA-Seq (**d**) densities between the two biological replicates. Read density is normalised in reads per million (RPM) of total reads, and is quantified either for individual ORFs (**c, d**) or individual codons (**e**).

**Figure 3.**
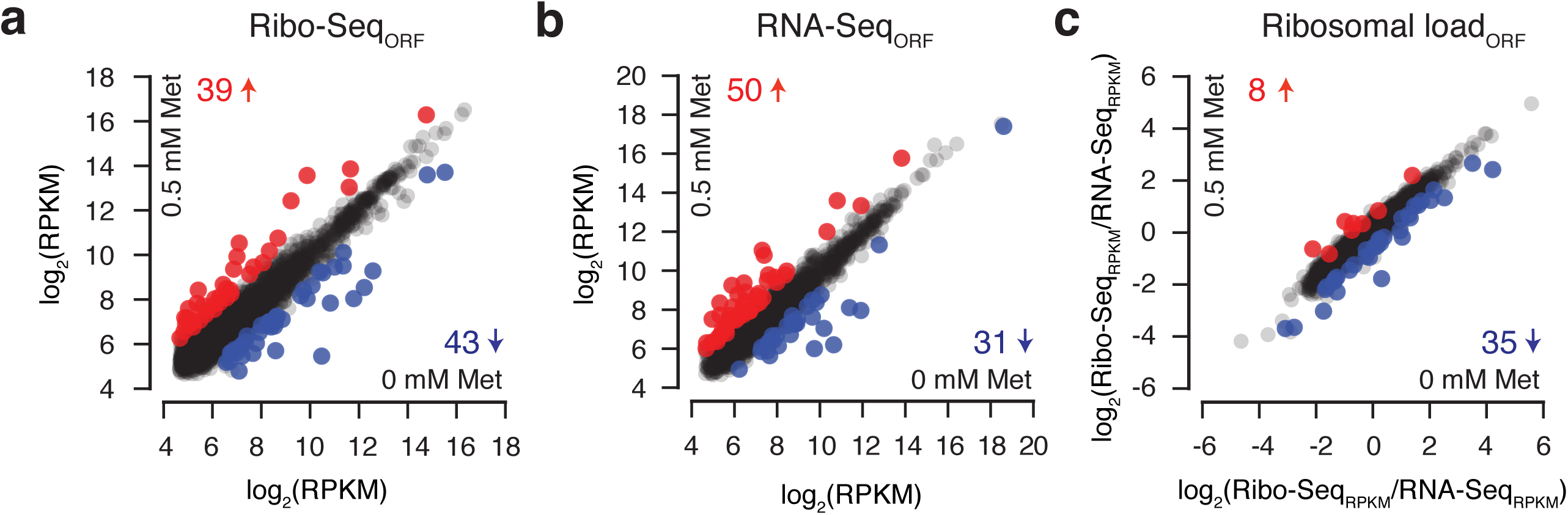
Genome-wide effects of eEF3 depletion on gene expression. The effects of eEF3 repression to ribosomal density (**a**), on the levels of mRNA (**b**) and ribosomal load (**c**) calculated for individual genes (2581 in total). Changes in gene coverage is considered significant when the Z-score of the differences is above 2 or below -2 in both replicates. Up-regulated genes in eEF3 depleted conditions are marked with red dots and down-regulated genes with blue dots, and the counts of genes presented as numbers in respective colors. Gene IDs of up-regulated and down-regulated genes, as well as associated GO IDs and p-values are provided in Supplementary Data S3. Read density is normalised as reads per kilobase per million (RPKM) of total reads for individual ORFs (**a** and **b**).

**Figure 4.**
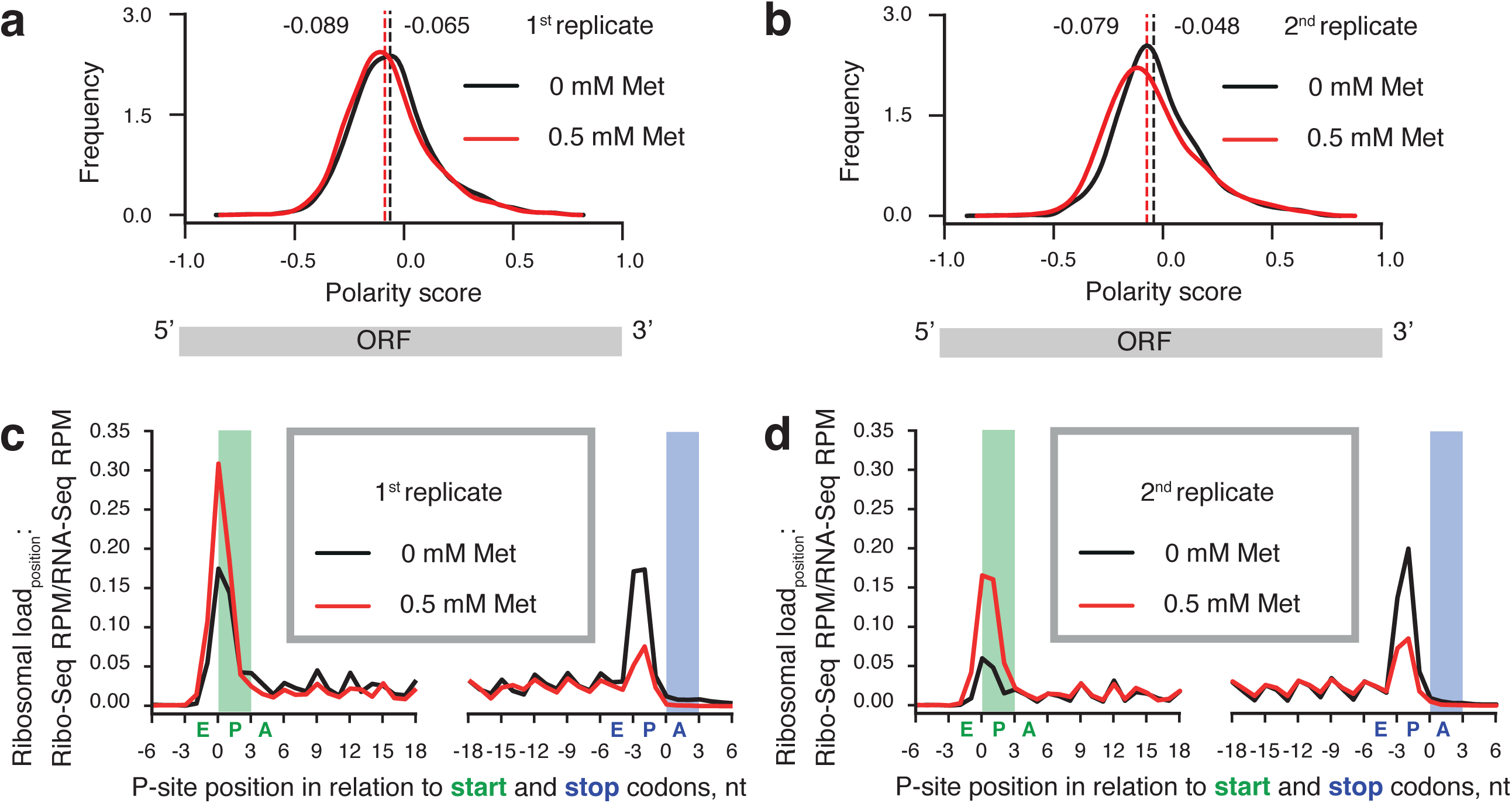
eEF3 depletion causes moderate redistribution of the ribosome density along mRNAs. The graphs show distributions of polarity scores for individual ORFs (**a** and **b**, two experimental replicates), and average ribosome density in the vicinity of start and stop codons (**c** and **d**, two experimental replicates). The expression of eEF3 was either repressed by the addition of 0.5 mM methionine (red trace) or unrepressed (black trace, no methionine). The locations of start and stop codons are highlighted with green and blue backdrops, and the corresponding A-, P- and E-site positions of initiating and terminating ribosomes are indicated with letters under the x-axis.

### Changes in gene expression upon eEF3 depletion are driven by metabolic adjustments and reduced growth rate

To uncover the effects of reduced abundance of eEF3 on gene expression we performed differential gene expression analysis and compared RNA-Seq_ORF_, Ribo-Seq_ORF_ and ribosomal load_ORF_ between the eEF3-deficient and eEF3-proficient cells. A considerable fraction of *S. cerevisiae* proteins is encoded by paralogous gene pairs, including highly expressed genes encoding ribosomal proteins and translation factors^26,27^. Therefore, if only single-mapping is allowed, a significant fraction of reads corresponding to these genes is lost (Supplementary Fig. S6). In order to avoid this systematic bias, for gene expression analysis we re-mapped the raw Ribo-Seq and RNA-Seq data retaining the reads that map twice. Differentially expressed (DE) genes were defined by Z-score <-2 or >2 of gene coverage difference between eEF3-deficient and eEF3-proficient conditions in both biological replicates (Supplementary Data S2). Analysis of the Ribo-Seq datasets identifies 82 genes that are differentially expressed in the eEF3-deficient condition compared to eEF3-proficient: 39 are up-regulated and 43 are down-regulated (Fig. 3a). The majority of the effect is driven by the mRNA copy number: 35 of the up-regulated genes are also up-regulated as per RNA-Seq analysis (i.e. 90% genes identified by Ribo-Seq) and 27 of the down-regulated genes are also picked up by RNA-Seq (63%) (Fig. 3b). Computing the ribosomal load subtracts the transcriptional effects to identify cases of specific translational regulation (Fig. 3c). The number of up-regulated genes is reduced to 8, consistent with predominantly transcriptional nature of the effect. However, out of 35 down-regulated genes only 4 are common between Ribo-Seq and ribosomal load datasets, indicating that this group of genes is predominantly down-regulated on the translational level.

We applied the YeastMine Gene Ontology (GO) enrichment tool^28^to identify the molecular functions, biological processes, and cellular components enriched in differentially expressed genes (Supplementary Data S3). GO annotations of genes with increased read density in Ribo-Seq and RNA-Seq point to biological processes related to the cell response to heat GO:0034605 (HSP104, DDR2, HSP12, HSP26, SSA4, TPS2) and stress GO:0070413 (HSP104, TPS2, TSL1). No specific GO enrichments were identified for the 8 genes that display specifically increased ribosome load. Induction of the heat shock response and elevated expression of chaperones upon eEF3 depletion could be a consequence of an accumulation of misfolded proteins due to defects in protein synthesis^29^. Genes with decreased RNA-Seq read density are dominated by GO categories related to metabolic processes, especially amino acid synthesis (GO:0017144, GO:0000096, GO:1901605, GO:0044281, GO:0006555 and GO:0006520; p-values from 4.33e^-6^ to 9.0e^-4^). This is an expected consequence of repressing eEF3 by the addition of 0.5 mM methionine to cell cultures. The gene set that displays a specific decrease in the ribosomal load_ORF_, shows a different pattern. The GO annotations are dominated by the cellular component categories: extracellular region, cell wall and cell surface (GO:0005576, GO:0009277, GO:0005618, GO:0030312 and GO:0009986; p-values from 5.3e^-5^ to 6.0e^-3^), most likely reflecting the significant growth rate decrease upon eEF3 depletion.

### eEF3 depletion causes a decrease in the efficiency of translation elongation

Defects in translation elongation lead to an accumulation of ribosomes on the 5’ part of ORFs^19,25,30^. This effect can be detected in ribosome profiling data by computing the so-called polarity score metric that ranges from -1 (corresponding to all of the ribosome density localised at the 5’ half of the ORF) to +1 (corresponding to all of the ribosome density localised at the 3’ half of the ORF)^30,31^. To account for potential differences in mRNA integrity, e.g. differential effects on co-translational mRNA degradation^32,33^, we normalised the Ribo-Seq reads to the RNA-Seq coverage for each individual nucleotide position (Ribosomal load_position_). Ribosome density corresponding to 15 nucleotides from both the 5’ and 3’ ends of ORFs were excluded from the analysis in order to avoid effects acting on initiation and termination, respectively. Consistent with a defect on translation elongation, depletion of eEF3 correlates with a slight but reproducible shift in ribosome distribution towards the 5’ end of ORFs (Fig. 4a and b). To test the statistical significance of the effect, we applied the nonparametric Wilcoxon signed-rank test of the null hypothesis i.e. that the two polarity score distributions originate from the same underlying distribution. In order to quantify biological variance we compared the two biological replicates. The p-values for eEF3-proficient (5.3e^-2^) and eEF3-deficient (5.5e^-4^) conditions provide an estimate of the biological variability. The difference between eEF3-proficient and eEF3-deficient conditions is significantly higher for both biological replicates (p-values 7.5e^-21^ and 2.1e^-21^). This suggests that the shift in ribosome distribution towards the 5’ end of ORFs is, indeed, specific to eEF3 depletion.

To determine the ribosomal load across the full length of all mRNAs, we performed a metagene analysis of ribosome density around the stop and start codons for all genes with sufficient coverage (Fig. 4c and d). Depletion of the ribosome recycling factor Rli1/ABCE1 leads to an increased ribosomal occupancy at stop codons and the appearance of aperiodic ribosome density in 3’-UTR regions^34^; if subunit dissociation during ribosome recycling is the primary function of eEF3^12^, we would expect a similar pattern. However, eEF3 depletion results in decreased ribosome occupancy at stop codons and does not lead to increased ribosome occupancy in the 3’-UTR regions, suggesting that eEF3 depletion compromises ribosome recycling less than it does elongation. Conversely, ribosome density increases at the start codon, suggesting that translation initiation becomes more rate-limiting upon eEF3 depletion.

### eEF3 depletion leads to a moderate decrease in ribosomal stalling on P-site proline residues

To uncover possible sequence-specific effects of eEF3 depletion we followed the approach originally developed by the Vázquez-Laslop and Mankin labs^35,36^. We computed changes in ribosome density at individual P-site codons between eEF3-deficient and eEF3-proficient cells – the relative fold difference, Ribo-Seq FD_P-site codon_ or just FD for simplicity (Fig. 5a). Positive log_2_(FD) values signify a relatively higher coverage of the individual codon by ribosomes under eEF3-deficient conditions, commonly interpreted as codon-specific ribosomal stalling – or at least slowing down or pausing of translating ribosomes^35,36^. Conversely, negative log_^2^_(FD) values indicate relatively lower ribosomal coverage under eEF3-deficient conditions. To discount the contribution of ribosome occupancy at initiation codons, the first 10 codons in all ORFs were excluded from the analysis. Assuming a near-normal distribution of the log_2_(FD), we identified codons with significantly decreased (Z-score <-2, i.e. the codon’s log_2_(FD) value is more than two standard deviations lower than the mean; 651 individual codons) or increased (Z-score >2; 856 individual codons) FD in both biological replicates (see Supplementary Data S4). For these codons we extracted the sequence starting from the A-site (+1 position), P-site (0 position), and E-site (-1 position), as well as for the positions of the amino acids in the polypeptide tunnel (from -2 to -5). To compute amino acid overrepresentation scores we used pLogo^37^that outputs log_10_-odds of over- or under-representation in relation to the background (see Supplementary Methods section in Supplementary information S1). The horizontal red bar in Fig. 5b-e represents the statistical significance threshold of the multiple test adjusted p-value (0.05).

**Figure 5.**
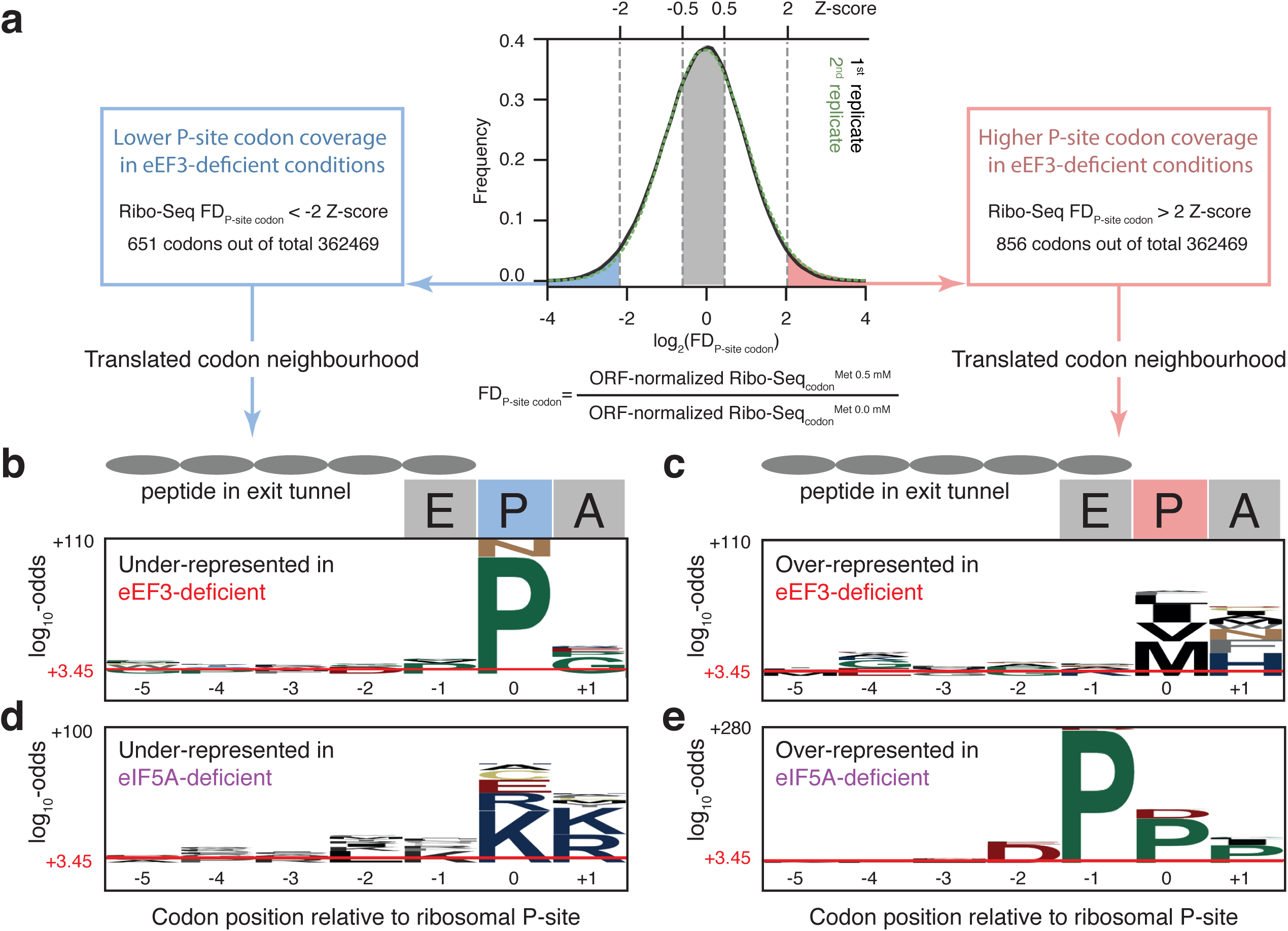
Amino acid- and position-specific redistribution of ribosomal density upon depletion of eEF3 or eIF5A. (**a**) Distribution of genomic sites (codons) according to the relative fold difference (FD) in the ribosome density between eEF3-deficient and eEF3-proficient cells. Positive FD values (right side of the distribution) indicate the sites where depletion of the factor leads to an increase in ribosome density, and negative FD values correspond to a relative decrease in ribosome density upon depletion of the factor. pLogo^37^was used to calculate overrepresentation of specific amino acids at positions relative to the P-site codon using codons with Z-score < -2 (eEF3: panel **b**, 651 codon positions; eIF5A^30^: panel **d**, 510 codon positions) and > 2 (eEF3: panel c, 856 codon positions; eIF5A^30^: panel **e**, 998 codon positions) that are common for both biological replicates. Horizontal red lines on the pLogos represent significance threshold (the log_10_-odds 3.45) corresponding to a Bonferroni corrected p-value of 0.05.

We detected no dramatic over-representation of specific amino acids in the E-site, indicating that the identity of the E-site tRNA does not influence the redistribution of ribosomal densities upon eEF3 depletion (Fig. 5a and c; Supplementary Table S2). This suggests that eEF3’s function in translation elongation is not E-site codon specific.

In the A-site, the strongest effect is overrepresentation of histidine-encoding codons amongst those with increased overage upon eEF3 depletion (Fig. 5c). Histidine is overrepresented 5.4 times (log_10_-odds of 19) in comparison to the background, followed by phenylalanine (2.6 times; log_10_-odds 9) and asparagine (2.5 times; log_10_-odds 6.6) in eEF3-deficient conditions. In the P-site, eEF3 deficiency leads to a moderate increase in ribosome density for hydrophobic amino acids, with the strongest signal being methionine (5 times; log^10^-odds 25) and valine (2.1 times; log_10_-odds 13).

The strongest amino acid-specific effect that we observe over all is a relative underrepresentation of proline codons in the P-site under eEF3-deficient conditions (Fig. 5b). That is, ribosomal density corresponding to P-site prolines is relatively higher in eEF3-proficient conditions. To put this effect into perspective, we re-analyzed the Ribo-Seq dataset of Schuller and colleagues who profiled a yeast strain depleted in the translation factor eIF5A^30^. Polyproline motifs are well-known to cause ribosomal pausing^38^, and eIF5A promotes translation of these stalling-inducing sequences^31,39^. Upon eIF5A depletion, ribosome pause sites (high FD; Z-score >2) are enriched with proline and aspartic acid (Fig. 5e; Supplementary Table S3). While proline codons are overrepresented in all of the three ribosomal sites, the E-site effect is the most dramatic (7.9 times; log_10_-odds 274). The next-strongest effect upon eIF5A depletion is overrepresentation of proline in the P-site (4 times; log_10_-odds 91). The strength of this effect is very similar to the proline overrepresentation in this position that we observe in eEF3-proficient conditions in relation to eEF3-depeleted (5.5 times; log_10_-odds of 93.5).

As in the case of eEF3, we detect a clear signal of relative underrepresentation of specific amino acids in the eIF5A-deficient strain, although the nature of amino acids is different. Positively charged amino acids lysine and arginine – well-known ribosomal stallers^40^-^42^ – are overrepresented 3.3-5.9 times in A- and P-sites (Figure 5d, Supplementary Figure 3; log_10_-odds of 13-39).

## Discussion

In this study, we have applied ribosome profiling to an eEF3-depleated *S. cerevisiae* strain under conditions of balanced growth. Surprisingly, the *depletion* of eEF3 leads to decreased occupancy of elongating ribosomes on P-site proline residues (Fig. 5b). The opposite effect, i.e. increased occupancy of elongating ribosomes on P-site proline residues is brought about by depletion of another E-site binder – translation factor eIF5A which promotes translation of polyprolines^30^ (Fig. 5e). Therefore, we hypothesise that lowering eEF3 concentration increases the E-site availability for eIF5A binding, increasing the efficiency of transpeptidation in the case of prolines. An alternative explanation is that the general reduction in the efficiency of translation elongation as observed in eEF3-deficient cells may simply mask the pauses that normally occur at proline codons, thus leading to an observed relative decrease in the ribosome density of P-site proline residues.

While eEF3 deficiency specifically *decreases* occupancy of elongating ribosomes on prolines, our re-analysis of the ribosome profiling dataset of the eIF5A-deficient strain^30^ has identified specifically *decreased* occupancy on lysine and arginine (Fig. 5d). It is instructive to contrast this result with that of Pelechano and Alepuz who performed 5Pseq, a genome-wide method of analysing translation by sequencing 5’-phosphorylated mRNA degradation intermediates^43^ This work used an eIF5A temperature-sensitive mutant^31^ rather than the degrone fusion used by Schuller and colleagues^30^. While Pelechano and Alepuz also detected lysine- and arginine-specific effects of eIF5A depletion, the effect is the opposite, i.e. lysine and arginine are overrepresented amongst the codons with increased ribosome occupancy in the eIF5A-depleted strain.

Since we detect no pile up of the ribosome protected fragments at the stop codon – a characteristic sign of defective ribosome recycling^34^– we conclude that ribosome recycling is not the primary function of eEF3. We also fail to detect any ribosomal stalling signature specific to the nature of E-site codon. From this we conclude that the identity of the E-site codon (and, hence the identity of deacylated E-site tRNA) does not seem to have a significant role in eEF3’s function as a general elongation factor.

## Methods

### Growth media, strains and genetic procedures

Yeast media were prepared as described^44^ with the difference that the composition of drop-out mix was as per Johansson^45^. Difco Yeast Nitrogen Base w/o Amino Acids was purchased from Becton Dickinson (291940), amino acids and other supplements from Sigma-Aldrich. YEPD medium supplemented with 200 µg/ml Geneticin^TM^ (Gibco 11811-023) was used to select for transformants containing the *kanMX6* marker^46^.

To construct a strain in which the expression of eEF3 (*YEF3*) is under control of the methionine repressible *MET25* promoter (P*_MET25_*), we first transformed a diploid strain, formed between BY4728 and BY4742^47^, with a *kanMX6*-P*_MET25_* DNA fragment with appropriate homologies. The *kanMX6*-P*_MET25_* DNA fragment was generated in three steps. The kanMX6 marker was amplified from pFA6a-*kanMX6* ^46^ using the primers V3 (5’-TATCCGGCCCCACCCATGCATAACCCTAAATTATTAGATCCGGATCCCCGGGTTAATTAA-3’), which introduces 40 bp homology to sequences ≈300 bp upstream of the *YEF3* ORF, and V4 (5’-GAATTCGAGCTCGTTTAAAC-3’). The P*_MET25_* fragment was amplified from genomic DNA from the BY4742 strain^47^using primers V5 (5’-GTTTAAACGAGCTCGAATTCGGATGCAAGGGTTCGAATC-3’), which introduces 20 bp homolgy to 3’ end of *kanMX6* fragment, and V6 (5’-GTTCTTCTAGAACCTTAATGGATTGCTGGGAATCAGACATTGTATGGATGGGGGTAA TAGA-3’), which introduces 40 bp homology to the 5’ end of *YEF3* open reading frame. The *kanMX6* and P*_MET25_* DNA fragments were then fused by overlap extension PCR^48^, generating the *kanMX6*-P*_MET25_* DNA fragment. After transformation, purifications by single cell streaks and PCR confirmation, a heterozygous P*_YEF3_*∷*kanMX6*-P*_MET25_*/*YEF3* strain was allowed to sporulate and the VKY8 (*MATα ura3Δ0 his3 leu2Δ0* P*_YEF3_*∷*kanMX6*-P*_MET25_*) and VKY9 (*MATα ura3Δ0 his3 leu2Δ0*) strains were obtained from a tetrad on SC-met-cys medium. The P*_YEF3_*∷*kanMX6*-P*_MET25_* allele in VKY8 was confirmed by PCR, using primers that annealed outside of sequences in the transformed DNA fragment, and subsequent DNA sequencing of the PCR product.

### Growth assays

The P*_MET25_*-*YEF3* (VKY8) strain was grown overnight in SC-met-cys medium at 30°C, diluted to an optical density at 600 nm (OD_600_) of 0.05 in SC-met-cys medium supplemented with different L-methionine concentrations. Cultures were grown at 30°C in a shaking water bath (New Brunswick™ Innova^®^ 3100) at 195 rpm. After 8 hours, the cultures were diluted to OD_600_≈0.05 in the same medium and grown overnight. In the morning, the cultures were re-diluted to OD_600_≈0.05 and growth was monitored through hourly OD_600_measurements. Growth rates (µ_2_) were calculated as slopes of linear regression lines through log_2_-transformed OD_600_data points.

The effect of the methionine concentration on the growth of wild-type (VKY9) and P*_MET25_*-*YEF3* (VKY8) cells on solid medium was determined from cultures grown overnight in liquid SC-met-cys medium. Cells were harvested, washed, serially diluted and spotted^49^ onto plates containing different methionine concentrations.

### Polysome profile analysis

The P*_MET25_*-*YEF3* (VKY8) strain was grown overnight in SC-met-cys medium at 30°C and diluted to an OD_600_of 0.05 in SC-met-cys and SC-cys (0.5 mM Met) medium. After 8h of growth at 30°C, the cultures were re-diluted in 150 ml of the same medium thereby ensuring that the OD_600_of the cultures was below 0.6 the next morning (15-17h). Growth was then monitored until the OD^600^ reached 0.8-1. A 50 ml aliquot of each culture was transferred into a pre-warmed flask and treated with 100 µg/ml cycloheximide, CHX, (C7698, Sigma-Aldrich) for 10 min under continuous shaking. Cells from both untreated and CHX treated cultures were pelleted by centrifugation at 3,000 x g for 5 minutes at room temperature, placed on ice, washed with 5 ml of ice cold Breaking buffer (20 mM Tris-HCl pH 7.4, 10 mM MgCl2, 100 mM KCl) with or without 100 µg/ml CHX, and pelleted again at 4°C. The pellet was resuspended in 250 µl of the respective Breaking buffer containing 1 mM DTT and 1x EDTA-free protease inhibitor cocktail and transferred to a 2 ml FastPrep-24 compatible microcentrifuge tube. Cells were lysed using 0.25 g of glass beads (0.5 mm diameter) and the FastPrep-24 for two 20 sec cycles at a speed setting of 4 m/sec with 1 min on ice between the steps. Lysates were cleared by centrifugation at max speed for 15 min at 4°C and 5 A_260_units loaded on precooled 7-45% linear sucrose gradients in SW41 tubes (made in Breaking buffer +/-CHX supplemented with 1 mM DTT and using a Biocomp Gradient Master instrument). Following centrifugation at 35,000 rpm for 3 hours at 4°C, the gradients were analyzed by measuring continuous absorbance at 260 nm using a Piston Gradient Fractionator (Biocomp Instruments).

### Preparation of NGS libraries (Ribo-Seq and RNA-Seq) and data analysis

The P*_MET25_*-*YEF3* (VKY8) strain was grown as described for the polysome profile analyses with the difference that the final culture volume was 750 ml and the cells harvested at OD_600_≈0.6 without prior CHX-treatment. Cells were harvested by rapid vacuum filtration onto a 0.45 µm nitrocellulose membrane, scraped off using a spatula, and frozen in liquid nitrogen. Cells were lysed by cryogenic milling in the presence of CHX-containing lysis buffer, and RNA-Seq and Ribo-Seq libraries were prepared from the cell extracts. RNA-Seq libraries were prepared using Scriptseq Complete Gold Yeast Kit (Epicentre). Ribo-Seq libraries were prepared essentially as per Ingolia and colleagues^18^with modifications on the rRNA removal and sample purification procedures. Detailed protocols can be found in our GitHub repository https://github.com/GCA-VH-lab/RiboSeqPy. Multiplexed Ribo-Seq and RNA-Seq libraries were sequenced for 51 cycles (single read) on an Illumina HiSeq2500 platform. Quality of Illumina reads was controlled using FastQC^50^, and low quality reads (Phred score below 20) were discarded. The adaptor sequence (5’-CTGTAGGCACCATCAAT-3’) was removed using Cutadapt^51^. After removing reads mapping to non-coding RNA, reads were mapped to *S. cerevisiae* reference genome R64-1-1.85 using HISAT2^52^. In the case of Ribo-Seq out of 132.5-143.5 million unprocessed reads, 46.7-68.9 million remained after removal of non-coding RNA and reads mapped multiple times. RNA-Seq reads were processed similarly, omitting the Cutadapt step; out of 78.5-87.3 million unprocessed reads, 51.2-59.0 million remained after removing non-coding RNA and reads mapped multiple times. Ribo-Seq data analysis was performed using custom software written in Python 3, available in GitHub at https://github.com/GCA-VH-lab/RiboSeqPy. For more detail see Supplementary information S1.

## Data availability

Ribo-Seq and RNA-Seq sequencing data have been deposited in the ArrayExpress database at EMBL-EBI (www.ebi.ac.uk/arrayexpress) under accession number E-MTAB-6938.

## Acknowledgements

We are grateful to Akira Kaji for sharing anti-eEF3 antibodies and Nicholas Ingolia for help sharing protocols for Ribo-Seq library preparation and help with data analysis. This work was supported by the funds from European Regional Development Fund through the Centre of Excellence for Molecular Cell Technology (VH); the Molecular Infection Medicine Sweden (MIMS) (VH); Swedish Research council (grant 2013-4680 to VH, 2015-04746 to GCA); Ragnar Söderberg foundation (VH); Magnus Bergvalls Foundation (2017-02098 to MJ); Åke Wibergs Foundation (M14-0207 to MJ); and Kempestiftelsernas grants (JCK-1627 to GCA and SMK-1349 to VH).

## Author contributions statement

VH conceived the study, coordinated the study, and drafted the manuscript together with VK, TM, GCA and MJ. VK, MJ and VH designed experiments and analyzed the data. TM and GCA analysed the ribosome profiling data. All authors have read and approved the manuscript as submitted.

